# Deep Modeling of Gain-of-Function Mutations on Androgen Receptor

**DOI:** 10.1101/2025.01.20.633961

**Authors:** Jiaying You, Jane Foo, Nada Lallous, Artem Cherkasov

## Abstract

The efficiency of Androgen Receptor (AR) pathway inhibitors for prostate cancer (PCa) is on decline due to resistance mechanisms including the occurrence of gain-of-function mutations on human androgen receptor (AR). Hence, understanding and predicting such mutations is crucial for developing effective PCa treatment strategies. Leveraging accumulated data on clinically relevant AR mutants with recent advances in deep modeling techniques, this study aims to unveil and quantify critical AR mutation-drug relationships. By incorporating molecular descriptors for drugs and mutated genes sequences, this work represented these features as single vectors and demonstrates their effectiveness in modeling AR mutant responses to conventional antiandrogens. The developed approach achieves up to 80% accuracy in predicting the gain-of-function behavior of AR mutants and therefore can potentially uncover unknown agonist/antagonist relationships among mutant-drug pairs.

## 1 Introduction

Prostate cancer is the second most common cancer in men, exerting a significant impact on their longevity and quality of life ^[1].^ The traditional treatment is by inhibiting the androgen receptor (AR) activity, which regulates biological responses in males, by inhibiting androgen production or interfering with their binding to the AR. However, confidence in AR-targeted drugs like enzalutamide ^[2]^ and abiraterone ^[3]^ is fading, as more patients exhibit adaptive resistance to these therapies.

Structurally, the AR gene is located on the X chromosome [4] and encodes three major domains: (1) the N terminal domain (NTD, residues 1-559), (2) the DNA binding domain (DBD, residues from 560-625), and (3) the C-terminal ligand binding domain (LBD, residues from 671 to 920). A flexible hinge region links the DBD to the LBD. Over the years, there has been tremendous progress in developing AR inhibitors with novel modalities [5] [6] [7]. However the LBD remains the only domain targeted by clinically approved antiandrogens such as darolutamide[8] and apalutamide[9], despite its susceptibility to resistance and gain-of-function mutations. To address this challenge, our lab has developed small molecules that disrupt AR function by targeting either the NTD [10] or the DBD [11]. Additionally, we are developing prediction models to assess the response of AR point mutation to inhibition by LBD-directed agents, employing machine learning techniques to capture agonist and antagonist behaviors within mutant-drug pairs.

In recent years, Quantitative Structure-Activity Relationship (QSAR) modeling has become as an effective method in bioinformatics, enabling researchers to elucidate the biological activity of chemical compounds based on their biomarkers such as molecular structures and protein sequences [12]. QSAR models can support various applications such as drug discovery [13] and toxicology prediction [14], by analyzing the interactions between biomarker descriptors and biological responses. The adoption of deep learning (DL)[15] models over traditionally used linear regression or statistical methods has significantly improved performance in this field, enabling the capture of complex patterns within high-dimensional data. DL models excel in automatically extracting informative embeddings from raw data inputs and learning through tailored iterations, making them well-suited for addressing many bioinformatics challenges, where biological datasets are often massive and biased. By leveraging neural networks, DL enhances the precision of QSAR models, allowing for higher accuracy in predicting compound activity, and empowering the exploration of novel molecular designs in large chemical spaces.

To provide a comprehensive understanding of how AR mutations impact responses to anti-androgen therapies and to foresee potential therapeutic strategies for patients with resistant prostate cancer, we incorporated data from previous research by Lallous et al. ^[16],^ which investigated mutant-drug interactions for 86 AR mutants across a panel of five anti-androgen drugs. Using this combined dataset, we successfully predicted AR mutant-drug interactions through machine learning technologies, including DL and traditional bench-marks. Using this combined dataset, we managed to predict AR mutant-drug interactions using machine learning technologies including DL and traditional benchmarks. Our experiment involved 4 major components: (1) dataset outlier detection, (2) feature generation and selection, (3) cross-validation, and (4) benchmark comparison. Through these steps, we established a robust classification model for AR mutant-drug pairs. This framework leverages machine learning techniques to model the complex relationships between AR mutations and their biological responses.

Our in-house dataset was generated from multiple experiments on 3 chemicals: enzalutamide, bicalutamide and darolutamide with 12 common mutants (Figure 1), where each AR-mutant pair was tested 8 times under 7 doses, were recorded the luciferase activities among all experiments and excluded values as outliers with 10% percentile (below the bottom 10% or above the top 90% percentile).

**Figure 1:**
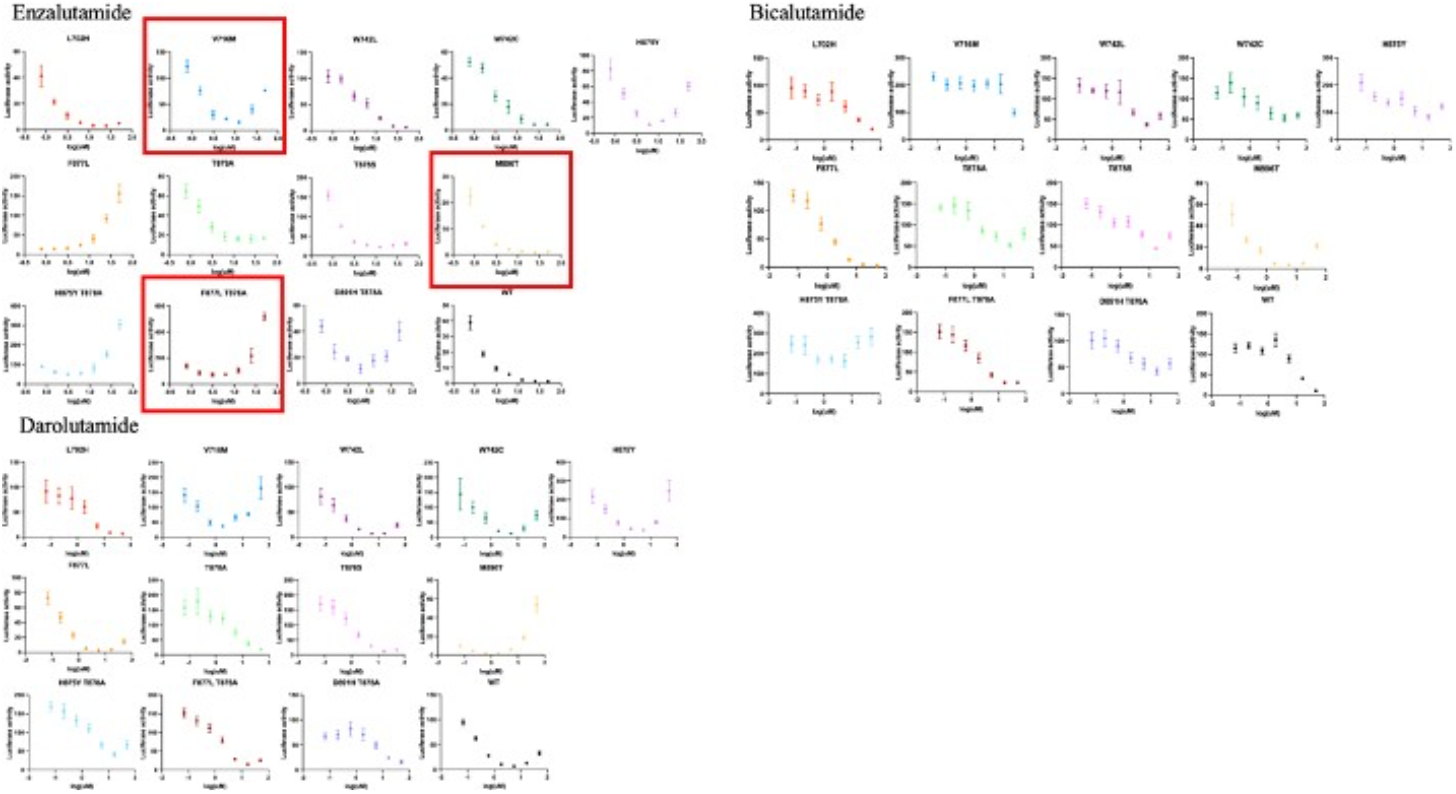
The effects of major antiandrogens were evaluated on 12 clinically observed resistant mutants. The graph features mutants’ responses to enzalutamide, bicalutamide and darolutamide. WT–the wild type, nonmutated AR. Red blocks are examples being labeled for up-(F877L_T878A), down-(M896T), mixed-(V716M) trend on enzalutamide.

Feature generation is also a critical step in machine learning modeling, here we utilized a variety of parameters including QSAR descriptors for drugs, and amino acid compositions for proteins, combined with physicochemical properties, to create a rich representation of mutant-drug compounds pairs. To retrieve robust prediction results, we used k-fold cross validation (k=3) when applying the DL model and our benchmark models including Support Vector Machine (SVM) ^[17]^ and Random Forest (RF)^[18]^, the model metric was then compared, we observed our DL model out-performs all traditional machine learning models.

In this work, we present a DL model designed to classify mutant-drug pairs based on their activity profiles. To evaluate its performance, we compared our deep neural network with established machine learning benchmarks, including random forest and support vector machines (SVM). Our aim is to leverage deep learning for multi-class classification tasks, enabling the accurate classification of drug responses in AR mutants.

## 2 Methods

### 2.1 Laboratory procedure for conducting a functional assay on PC3 cells

#### 2.1.1 Cell lines

PC3 cells were purchased from the American Type Culture Collection (ATCC) and were maintained in RPMI 1640 (Gibco, Life Technologies) supplemented with 10% fetal bovine serum (FBS). The cells were cultured in a humidified incubator at 37°C and 5% CO2 and routinely checked for mycoplasma contamination.

#### 2.1.2 Luciferase transcriptional assay

PC3 cells were starved for 72 hours in RPMI 1640 supplemented with charcoal-stripped FBS (CSS) then seeded into a 96-well plate at a density of 5000 cells/well. The following day, the cells were transfected with pcDNA-AR (wildtype or mutant) and ARR3tk-luc luciferase reporter plasmid ^[19]^ 24 hours post transfection, the cells were stimulated with 0.1nM R1881 and treated with a 1:3 serial dilution from 50 *µ*M to 0.07 *µ*M of enzalutamide simultaneously. After 24 hours, the cells were lysed with 50 *µ*l of passive lysis buffer (Promega #E1910) and 20 *µ*l of lysate was transferred to a white flat-bottom 96-well plate (Corning Life Sciences Cat#3912). Luciferase activity was measured using luminescence readings after adding 50 *µ*l of luciferase reagent (Promega Luciferase Assay System #E1500). Luciferase activity was normalized and expressed as a percentage of wildtype AR activity.

### 2.2 Dataset generation

In this work we additionally considered 12 clinically relevant AR mutants, which were experimentally tested against 3 known AR antiandrogens: enzalutamide, bicalutamide and darolutamide. We recorded their responses by measuring the luciferase-labeled transcriptional activities. The assay involves transfecting the cells with wild-type or mutated AR, treating them with compounds of interest, and measuring luminescence as an indicator of AR transcription. The results are then normalized and expressed in percentage of wild-type AR activity. We labelled the dose-response reaction amino acid substitutions of AR by common anti-androgen drugs into 3 categories, antagonist agonist and mixed, where we observe a up/mixed/down pattern correspondingly shown in red blocks in Figure 1 on various mutants treated with enzalutamide.

To enrich our dataset and enhance the depth of modeling, we augmented the data by incorporating findings from the previous work ^[20]^ on AR mutants. In particular, we integrated data from an experiment investigating the reactions of 86 AR mutants to 5 anti-androgen chemicals: bicalutamide, enzalutamide, hydroxyflutamide, apalutamide and darolutamide. This dataset was originally published by Lallous et al. Combined with the experimental data for 12 additional mutants from this study, we labelled the resulting combined datasets with the following rules:

1. A repeated of eight experiments were conducted at each concentration for a single mutant-drug pair, we first calculated the modified z-score, and removed data points from repeated experiments with outliers, threshold for this step is two standard deviations.
2. Average of luciferase activity scores were then compared on different concentration levels from -2 to 2 (in log form, unit: *µ*M). If the luciferase activation values only increase with higher concentration, we label such pairs as class 1 (up pattern class). If luciferase activation values only decrease with higher concentration, we label such pairs as class 2 (down pattern class). If we observe a mixed pattern of ups and downs on various concentration values, we label those pairs as class 3(mixed pattern class). Note that if the luciferase activation values are stable with all drug doses, we included those pairs in the mixed pattern class as well. These labeling are later being used in multi-class classification modeling.
3. A combination of the above dataset was generated eventually, including five unique anti-androgen compounds and 86 unique mutants. 238 mutant-drug pairs were labelled for either down, up or mixed (no response).

### 2.3 Feature generation

Feature representations are critical in deep modelling and the common feature generation technologies include one-hot encoding ^[21]^, image feature extraction^[22]^, time series modeling^[23]^, data normalization and scaling^[24]^. The objective of finding appropriate feature is to transfer data points into the format that can be comprehended to neural networks.

In artificial intelligence, feature vectors formed the input layer, the starting point where massive neuros in hidden layers can be learned and progressed actively during the intensive computational steps. Sequence embedding is one of the most common and effective encoding for sequence data point such as genes ^[25].^ In our study, we focused on mutated sequence data and encoded the mutants with descriptors. Those descriptors include:

#### QSAR descriptors

We explored molecular parameters including BLOSUM indices ^[26],^ Cruciani properties ^[25]^, FASGAI vectors^[27]^, Kidera factors^[28]^, MS-WHIM scores^[29]^, ProtFP descriptors^[30]^, ST-scales^[31]^, T-scales^[32]^, VHSE-scales^[26]^, IND descriptors^[33] [34] [35]^ and Z-scales^[18]^. Those QSAR descriptors capture various physicochemical and structural properties of molecules. These descriptors can be useful for characterizing the chemical and structural changes induced by mutations and predicting their effects on protein function or interaction with other molecules.

#### Amino acid counts and frequency

Analyzing the counts and frequencies of different amino acids in mutant sequences can provide insights into the composition and distribution of amino acids, which could be important for understanding the impact of mutations on protein structure and function.

#### Sequence profiles

These included hydrophobicity, hydrophobic moment, and membrane position, provide information about the local environment and structural features of amino acids within a protein sequence.

#### Physicochemical Properties

Physicochemical properties like the aliphatic index, instability index, theoretical net charge, isoelectric point, and molecular weight provide additional insights into the biochemical characteristics of mutant proteins. These properties can influence protein stability, solubility, and interaction with other molecules, making them relevant for understanding the functional consequences of mutations.

#### Biological Properties

Predicting structural class and other biological properties based on mutant sequences can provide valuable information about the potential functional impact of mutations. This could include predictions of protein secondary structure, subcellular localization, or functional domains.

#### Morgan fingerprints

These fingerprints generally used in machine learning models for molecular embeddings in various domain such as drug-drug interaction ^[36]^ prediction, drug-target interaction ^[37]^ prediction and drug repurposing ^[38]^. Morgan fingerprints is considered a balance between simplicity, efficiency, and representational power, making them one of the most widely used fingerprinting methods in cheminformatics. In this work, we extracted drug descriptors from Simplified Molecular Input Line Entry System (SMILE) with python package RDKit (RDKit: Open-source cheminformatics; http://www.rdkit.org), and generated morgan fingerprints with radius = 2, meaning with 2 diameters of the atom envi ronment are considered for encoding. We generated 128 bits of morgan fingerprints as drug descriptors embedded with binary 0/1 vectors.

We later combined heterogenous encodings of mutants and drugs into single vectors and fed them into the deep neural network for deep modeling and other traditional machine learning benchmark models.

### 2.4 Deep modeling

Deep learning is an artificial intelligence technology that learns data points through a bunch of neurons and layers actively by forward and backward propagations. The weighs assigned to each neural are assigned initially with non-sense numbers as a starting point, and graduate begin to update towards one goal: minimize the difference between real target value and predicted target value by the learned linear or unilinear formula. It has been widely applied into various fields such as image detection ^[39]^, language recognition^[40]^, drug discovery^[38]^ and recommendation systems^[41]^. The power of deep learning is the ability to learn automatically like human being with hierarchical feature representations. Compared with traditional machine learning technologies, deep learning can generally take massive data points and process them in an un-linear pattern, making it effective for complex concepts where intricate correlations may not be picked up by tradition linear models (Figure 2).

**Figure 2:**
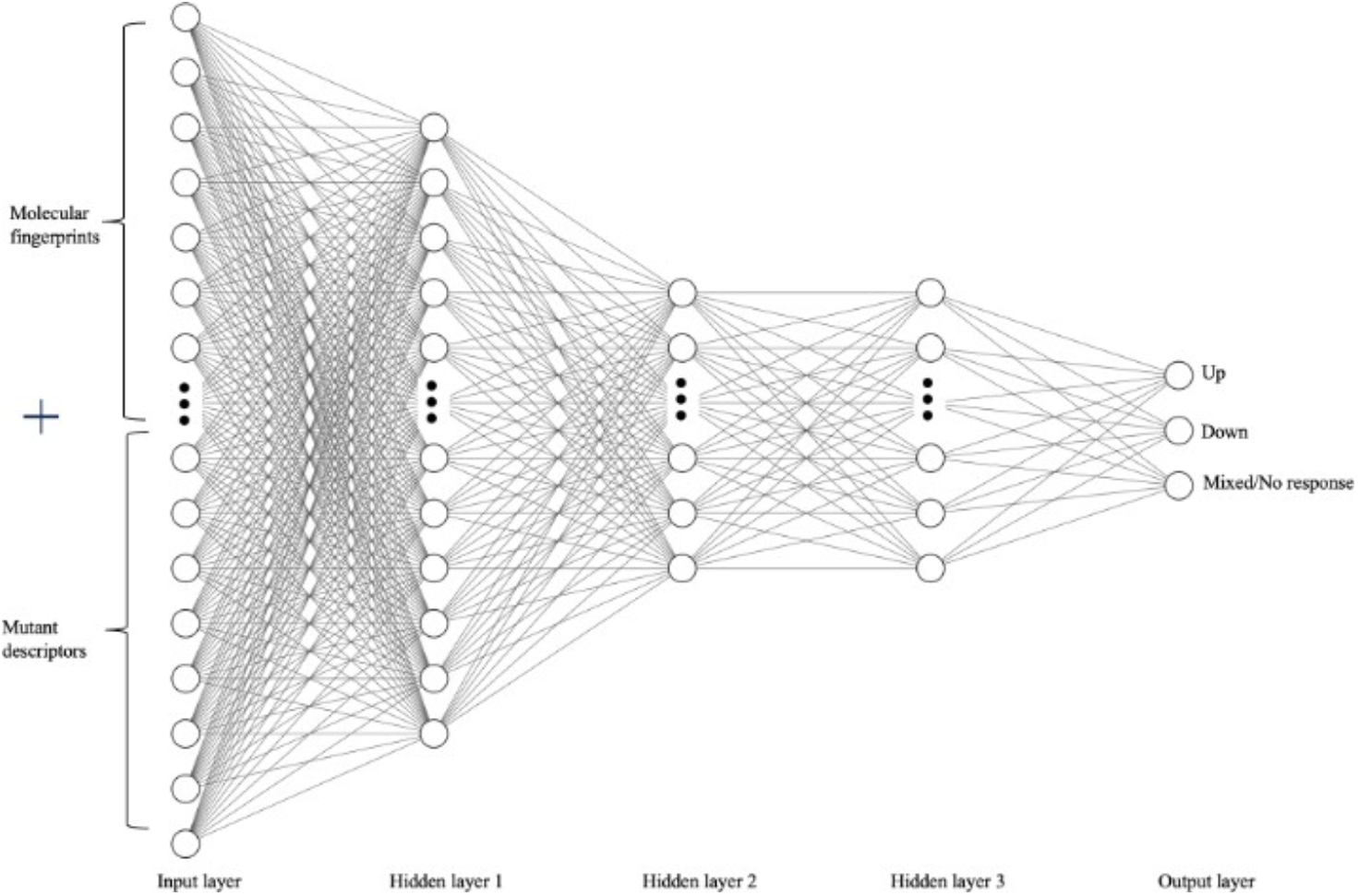
Deep neural network structure. Usually contains an input layer with sample representations, hidden layers with neurons that learns weighs in training process, and one output layer for classification responses.

In this work, we proposed a deep neural network model composing three hidden layers with 64, 32, 32 neurons respectively for multi-class classification. Heterogenous descriptors on mutants and drugs are combined as the input layer, and the AR activity on mutant-drug pairs were labeled as agonist, antagonist and mixed responses as the final output layer.

### 2.5 Benchmark modeling

To validate the performance of our deep neural network (DNN) model, it is crucial to select benchmark models that are widely acknowledged in the field for comparison. Among these, state-of-the-art machine learning algorithms such as Random Forest and Support Vector Machines (SVM) are particularly noteworthy for their effectiveness in classifying data points into three or more categories.

#### 2.5.1 Random Forest

Random forest in an ensemble learning method that binds multiple decision trees for training and votes their outputs for final predictions. Using randomly selected sub features, each tree in the random forest system makes a prediction independently, the randomness enables the random forest to capture various patterns within the data. This method is ideal as our benchmark model for predicting mutants-drug responses, as it classifies multi response types by dividing the feature space into different classes. It is particularly effective handling complex relationships with large variables data points by enhancing robustness and preventing overfitting, given its assembling nature of having numerous for independent predictions.

#### 2.5.2 Support Vector Machines (SVM)

SVM is a supervised learning model that classifies datapoints by optimizing a hyperplane (N dimensional space) that maximizes the distance between different classes. Those closest data samples are called support vectors for each class, and the goal is to find the decision boundary that maximize the margin between those support vectors. The decision boundary will be a straight line in binary classification problems if a linear relationship is caught between labels and features, and a higher dimensional hyperplane will be utilized when datapoints are not linearly separable. To handle the non-linearity of datapoints. Here we applied non-linear SVM that utilizes kernel function and brings dataset to a higher dimension, where again a straight line can be achieved separating different classes linearly.

In this paper, we measured the performance using Random Forest and SVM on mutant-drug pairs using identical embedded molecular descriptors. By comparing their performance metrics obtained from our deep learning model and benchmark models, we aim to gain insights into the strengths and limitations of each approach, highlighting the advantages of deep learning in this specific application.

## 3 Results

### 3.1 Data generation

In this study, we constructed the experiment that records the drug responses of various androgen receptor (AR) mutants to anti-androgen drugs enzalutamide, bicalutamide, darolutamide, hydroxyflutamide, apalutamide. Each of the experiment contains unique AR mutants, each tested by a single drug under seven different concentrations.

To ensure the integrity and reliability of our dataset, we applied a multi-step labeling process:

#### Outlier Removal

Each experimental result detected for outlies before labelling. We identified extreme values in data points that fell outside the 10th and 90th percentiles.

#### Average Calculation

After outlier removal, we calculated the average luciferase activities for each mutant-drug pair across the different concentrations.

#### Trend Classification

##### Upward Trend

Labeled as “up” if luciferase activity consistently increased with higher concentrations.

##### Downward Trend

Labeled as “down” if activity consistently decreased with higher concentrations.

##### Mixed Trend

Labeled as “mixed” if the response exhibited both increases and decreases or no response across different concentrations.

This systematic approach resulted in a dataset containing nearly 240 mutant-drug pairs, each classified into one of the three categories (up, down, or mixed). We visualized the statistics on interaction pairs on 5 unique drugs (Figure 3) and 86 unique mutants (Figure 4) respectively in below graphs.

**Figure 3:**
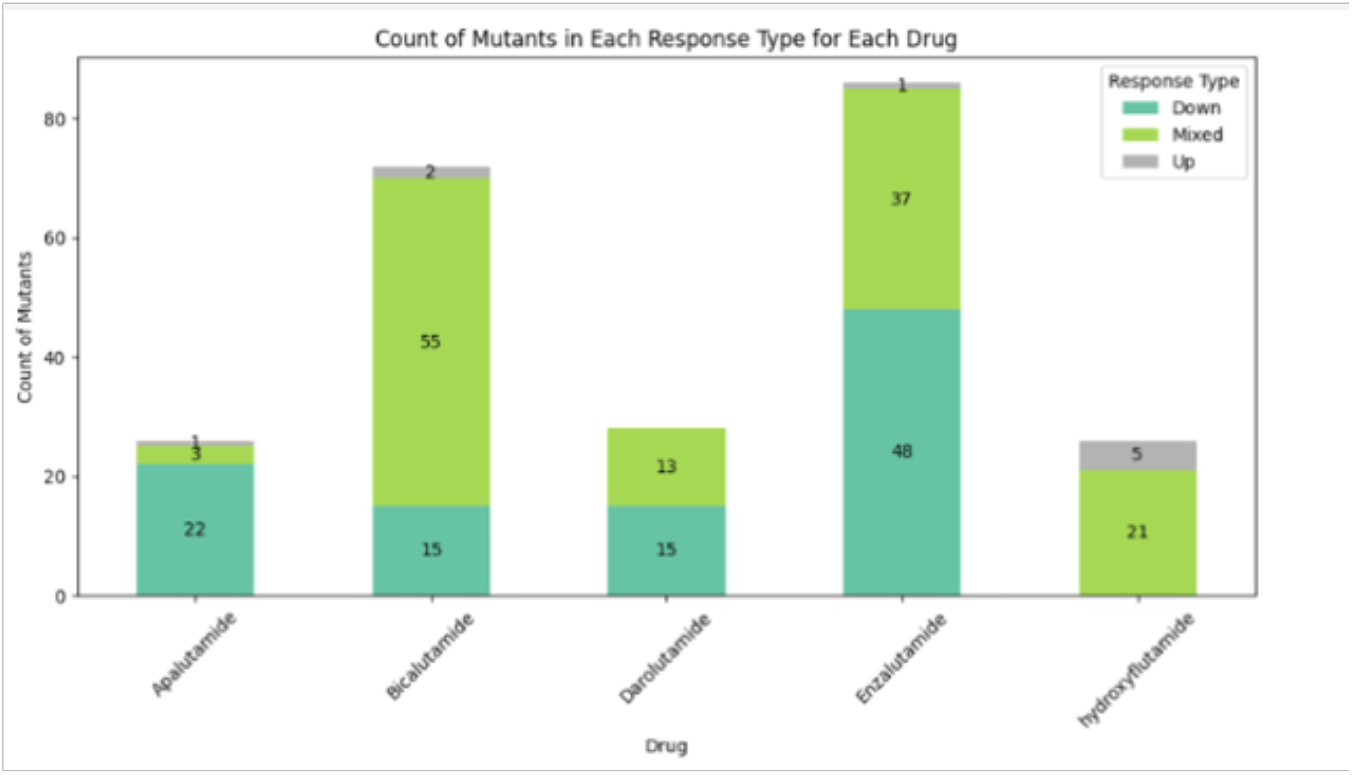
Count of mutant-drug response types on 5 antiandrogen drugs: enzalutamide, bicalutamide, darolutamide, hydroxyflutamide, apalutamide in the combined dataset

**Figure 4:**
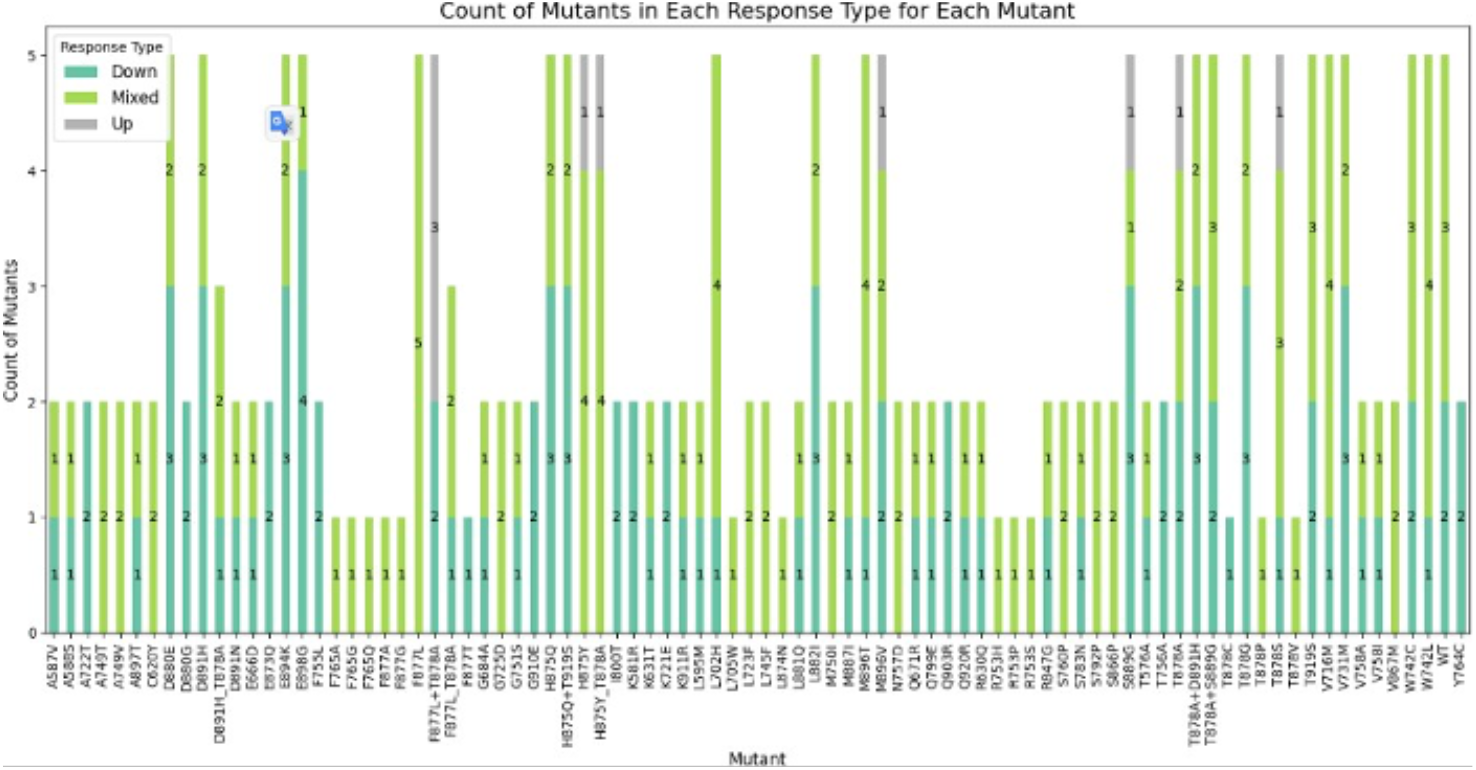
Count of mutant-drug response types on 65 unique mutants in dataset

### 3.2 Deep learning model performance

We conducted model training through a systematic approach to predict the responses of androgen receptor mutants to anti-androgen drugs. Initially, we merged comprehensive features in one vector including drug embedding and mutant descriptors. The response labels were encoded into numeric classes. Feature dimensionality was reduced using Principal Component Analysis (PCA)^[42]^ analysis, retaining 64 principal components before feeding into the neural network model. This optimized feature set was subsequently split using shuffle split 3-fold cross-validation, ensuring robust training and evaluation with 70% of the data for training and 30% for testing.

A feedforward neural network was conducted using PyTorch with 3 fully connected hidden layers with ReLU^[43]^ activations that provides non-linear calculations, and a sigmoid output layer that transfers output into possibilities. The model was trained using binary cross-entropy loss and stochastic gradient descent for optimization. Performance metrics including accuracy, precision, recall, and AUC-ROC were monitored to evaluate the model’s effectiveness in below figure 6.

Finally, the trained model’s predictions were compiled for all mutant-drug combinations, enabling the identification of potential new drug candidates for experimental validation. The DNN model attained an accuracy score of 0.86, an AUC values above 0.91 (Figure 5) and the lowest precision score of 0.8. We also plotted the training losses and accuracy over epochs in visual to prevent over fitting or under fitting shown in figure 6.

**Figure 5:**
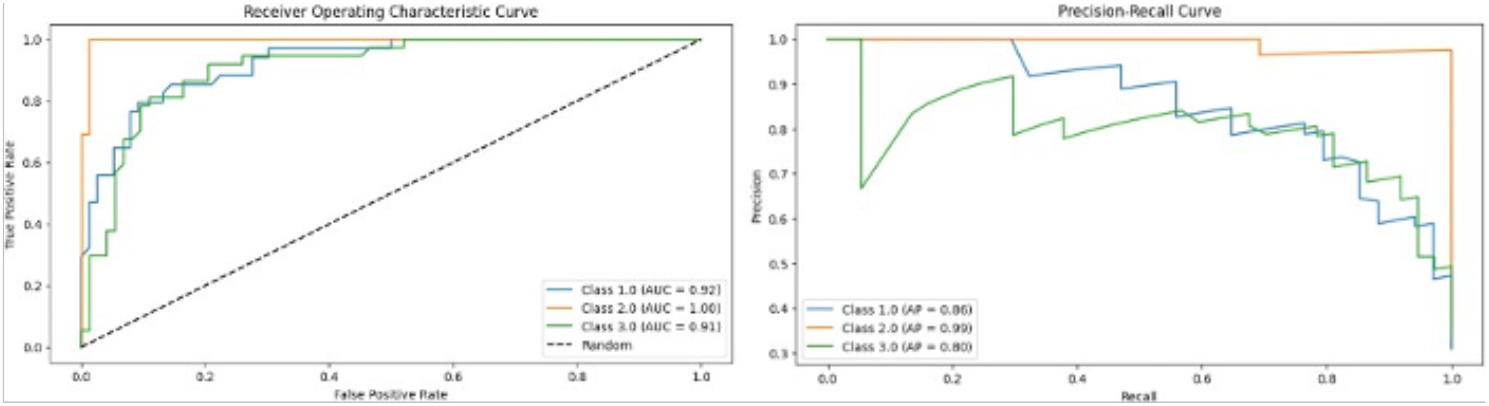
DNN model performance in accuracy, AUCand precision. The model accuracy is 0.85 and the AUC values above 0.89 wasachieved

**Figure 6:**
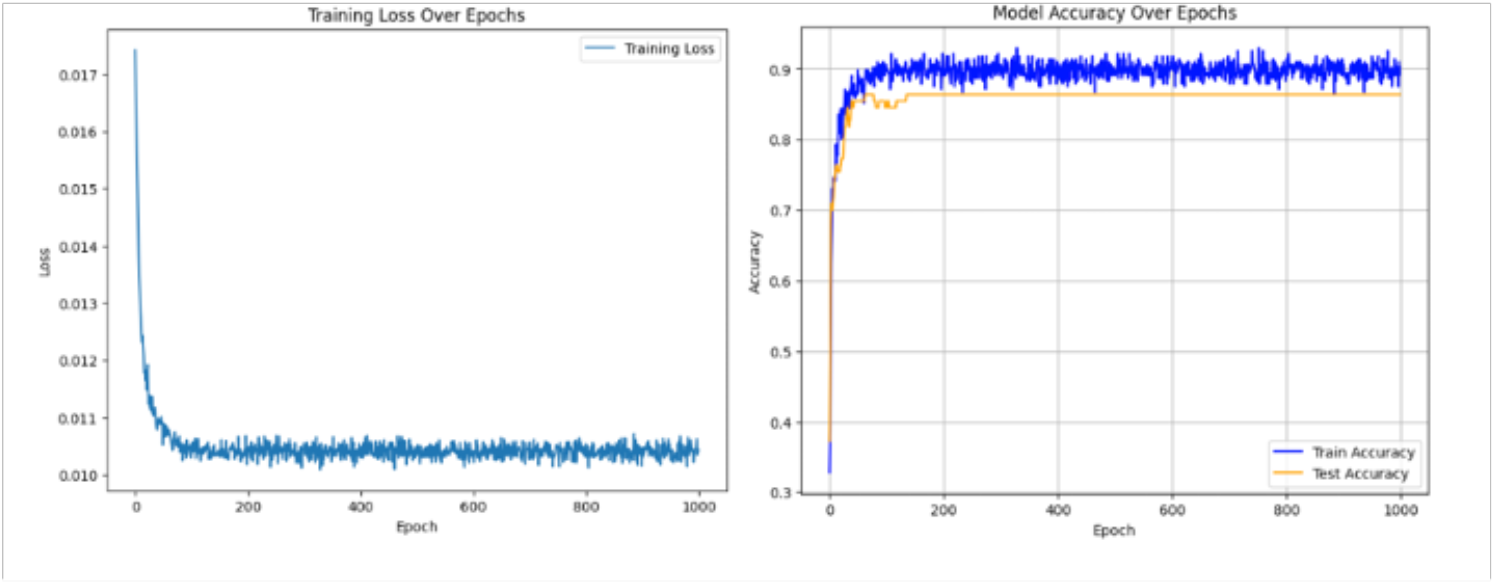
Training loss decreases, and recall/accuracy increases in training iterations.

### 3.3 Benchmark model performance

To evaluate the performance of the predictive models, we employed two widely used machine learning algorithms: Support Vector Machine (SVM) and Random Forest (RF). These models were selected for their robustness in handling multi-class classification tasks and their ability to provide interpretable results.

We first implemented Support Vector Machine using Scikit-learn library in python, and enabled probability in returns for AUC calculation. To be consistent with the primary neural work model the SVM was trained on the same integrated dataset consisting of feature representations generated from mutant and drug descriptors. The performance of the SVM model was evaluated an overall accuracy of 83%. The Random Forest model was constructed using the Scikit-learn library again using the same feature set generated from mutant-drug pairs. The RF model’s performance was also evaluated, yielding an overall accuracy of 84%.

The performance of benchmark classification models was evaluated using the Area Under the Curve (AUC) of the Receiver Operating Characteristic (ROC) curve to compare with the primary neural network model. Figure 7 below presents the ROC curves for both the Random Forest and Support Vector Machine (SVM), where the false positive rate (FPR) is displayed on the x-axis, and the true positive rate (TPR) on the y-axis. The ROC curve illustrates the trade-off between sensitivity and specificity at different threshold settings. The diagonal dashed line represents the performance of a random classifier as a baseline for comparison. The AUC values for the models were calculated from the ROC curves for both models. The Random Forest model achieved an AUC of approximately 0.87, indicating its strong capability to discriminate between classes. The SVM model displayed an AUC of approximately 0.90, indicating a better prediction power compared with random forest model. These values suggest that both models exhibit valuable predictive performance, with the SVM model marginally outperforming the random forest in terms of area ROC value shown in below figure 7.

**Figure 7:**
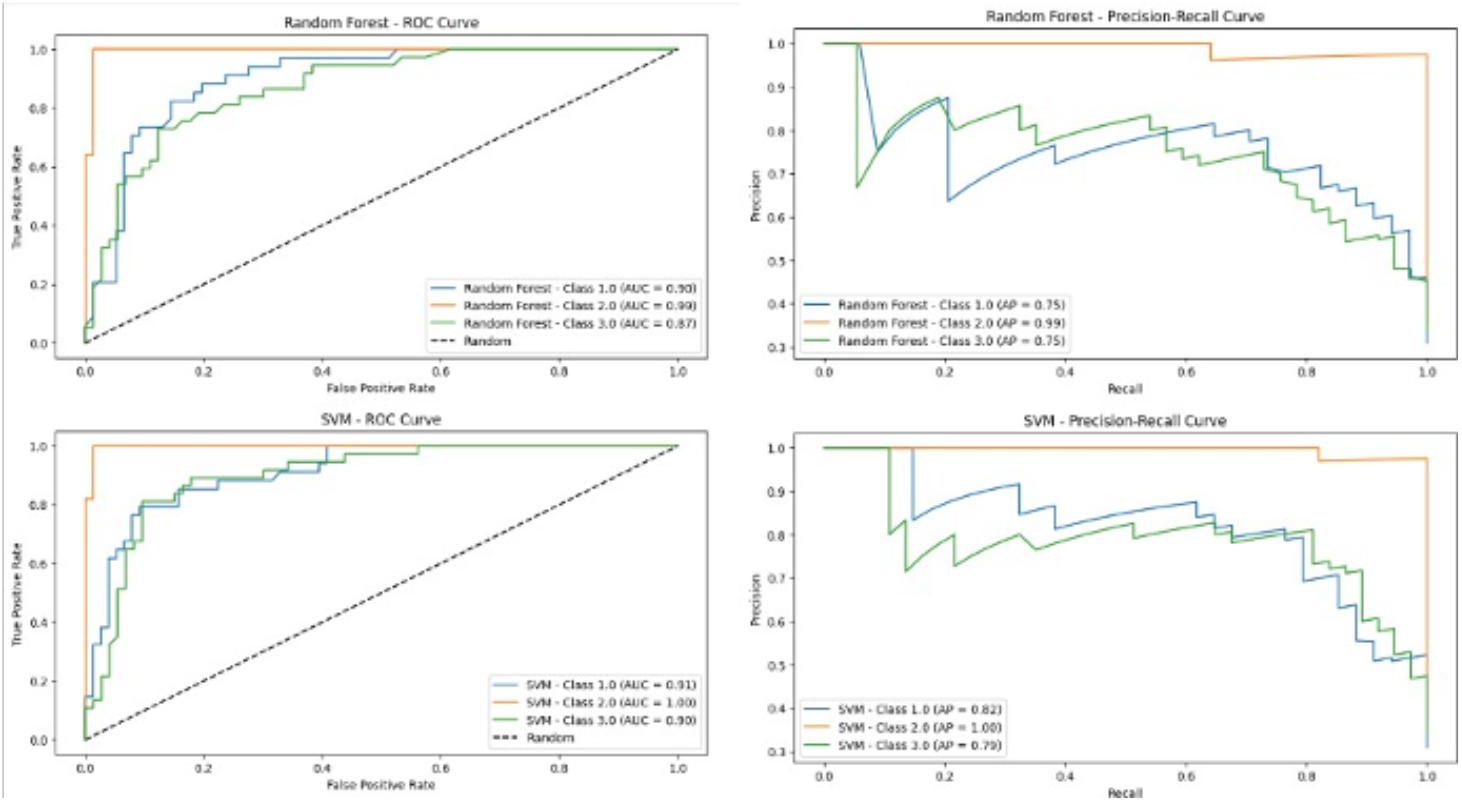
ROC and precision-recall curves for two benchmark models: Random Forest and SVM.

## 4 Discussion

Our research shows that by using DL models with integrated embeddings from mutant and drug descriptors, we can achieve outstanding performance in predicting AR mutants’ responses towards anti-androgen therapies. While people are losing confidence in traditional treatments due to gain-of-function mutations in the AR, machine learning model shows the potential in foreseeing drug responses that can be utilized in novel drug design or distinguishing possible novel mutants. In this paper, we capitalized on the integration of molecular descriptors and gene sequence data as informative representation and fed them in the unified format to the neural network model. By leveraging the comprehensive dataset of AR mutants and a diverse range of anti-androgen compounds, we were able to achieve an 79% accuracy in mutant-drug pair predictions, underscoring the potential of machine learning techniques in this domain. This approach allows for the exploration of previously insufficiently studied interactions and potentially unveiling new therapeutic strategies. The successful classification of mutant-drug pairs into distinct response categories—agonist, antagonist, and mixed— demonstrates the robustness of our model in capturing the various effects on mutations.

Benchmark models including random forest and support vector machine were established for comparison. We calculated the accuracy and plotted their ROC curves to demonstrate their prediction power. Tho the random forest and SVM also shows the abilities in distinguishing drug responses in multi-class on AR mutants, they both underperformed compared with the developed neural network model, which exhibits superior capabilities in capturing non-linear patterns and achieving higher accuracy. These findings demonstrate the potential in deep learning for mutant-drug response prediction and its capacity to facilitate novel medicine therapies for prostate cancer patients.

Moreover, to ensure the reliability and integrity of this research, the multi-step cleaning was applied in data generation and labeling process in this study. The processed dataset containing a diverse response of nearly 70 AR mutants and 5 known anti-androgen drugs, becomes a significant resource that can support future investigations into AR biology and therapeutic resistance.

## 5 Conclusion

This paper investigated the efficiency of deep learning models in predicting the agonist/antagonist behavior within AR mutant-drug pairs. The approach relied on the use of combined embeddings on sequence- and molecular QSAR descriptors and enabled high accuracy predictions. This study not only demonstrates potentials in predicting AR mutations-drug responses using deep learning but also enables effective exploration of novel antiandrogens with reduced or eliminated ability active AR mutants. It is anticipated that the developed approach will help tackling clinical challenges caused by AR mutations and ultimately improve prostate cancer patient care.

## 6 Supplementary information

Supplementary data are available online at https://github.com/chill-bear/ar_mutants/tree/add_files.

## Notes

### Competing Interest Statement

The authors have declared no competing interest.

